# Autoantibody landscape of advanced prostate cancer

**DOI:** 10.1101/2020.05.02.074575

**Authors:** William S. Chen, Winston A. Haynes, Rebecca Waitz, Kathy Kamath, Agustin Vega-Crespo, Raunak Shrestha, Minlu Zhang, Adam Foye, Ignacio Baselga Carretero, Ivan Garcilazo Perez, Meng Zhang, Shuang G. Zhao, Martin Sjöström, David A. Quigley, Jonathan Chou, Tomasz M. Beer, Matthew Rettig, Martin Gleave, Christopher P. Evans, Primo Lara, Kim N. Chi, Robert E. Reiter, Joshi J. Alumkal, Rahul Aggarwal, Eric J. Small, Patrick S. Daugherty, Antoni Ribas, David Y. Oh, John C. Shon, Felix Y. Feng

**Affiliations:** Department of Radiation Oncology, University of California, San Francisco, San Francisco, CA, United States; Helen Diller Family Comprehensive Cancer Center, University of California, San Francisco, San Francisco, CA, United States; Serimmune, Inc. Santa Barbara, CA, United States; Division of Hematology and Oncology, University of California, Los Angeles, Los Angeles, CA, United States; Department of Medicine, University of California, San Francisco, San Francisco, CA, United States; Knight Cancer Institute, Oregon Health & Science University, Portland, Oregon, United States; VA Greater Los Angeles Healthcare System, Los Angeles, CA, United States; University of British Columbia, Vancouver, British Columbia, Canada; University of California Davis, Davis, CA, USA; Department of Urology, University of California, Los Angeles, Los Angeles, CA, United States; Department of Hematology and Oncology, University of Michigan, Ann Arbor, MI, United States; Department of Urology, University of California, San Francisco, San Francisco, CA, United States

## Abstract

Although the importance of T-cell immune responses is well appreciated in cancer, autoantibody responses are less well-characterized. Nevertheless, autoantibody responses are of great interest, as they may be concordant with T-cell responses to cancer antigens or predictive of response to cancer immunotherapies. We performed serum epitope repertoire analysis (SERA) on a total of 1,229 serum samples obtained from a cohort of 72 men with metastatic castration-resistant prostate cancer (mCRPC) and 1,157 healthy control patients to characterize the autoantibody landscape of mCRPC. Using whole-genome sequencing results from paired solid-tumor metastasis biopsies and germline specimens, we identified tumor-specific epitopes in 29 mutant and 11 non-mutant proteins. Autoantibody enrichments for the top candidate autoantigen (NY-ESO-1) were validated using ELISA performed on the prostate cancer cohort and an independent cohort of 106 patients with melanoma. Our study recovers antigens of known importance and identifies novel tumor-specific epitopes of translational interest in advanced prostate cancer.

**Statement of significance:** Autoantibodies have been shown to inform treatment response and candidate drug targets in various cancers. We present the first large-scale profiling of autoantibodies in advanced prostate cancer, utilizing a new next-generation sequencing-based approach to antibody profiling to reveal novel cancer-specific antigens and epitopes.

**Disclosure of Potential Conflicts of Interest:** JJA reports receiving consulting income from Janssen Biotech and Merck and honoraria from Astellas for speaker’s fees. MR reports receiving commercial research support from Novartis, Johnson & Johnson, Merck, Astellas, and Medivation, and is a consultant/advisory board member for Constellation Pharmaceuticals, Amgen, Ambrx, Johnson & Johnson, and Bayer. A.R. has received honoraria from consulting with Amgen, Bristol-Myers Squibb, Chugai, Dynavax, Genentech, Merck, Nektar, Novartis, Roche and Sanofi, is or has been a member of the scientific advisory board and holds stock in Advaxis, Arcus Biosciences, Bioncotech Therapeutics, Compugen, CytomX, Five Prime, RAPT, ImaginAb, Isoplexis, Kite-Gilead, Lutris Pharma, Merus, PACT Pharma, Rgenix and Tango Therapeutics. FYF serves on the advisory board for Dendreon, EMD Serono, Janssen Oncology, Ferring, Sanofi, Blue Earth Diagnostics, Celgene, consults for Bayer, Medivation/Astellas, Genetech, and Nutcracker Therapeutics, has honoraria from Clovis Oncology, and is a founder and has an ownership stake in PFS Genomics. SGZ and FYF have patent applications with Decipher Biosciences. SGZ and FYF have a patent application licensed to PFS Genomics. SGZ and FYF have patent applications with Celgene. WAH, RW, KK, PSD, and JCS have ownership of stocks or shares at Serimmune, paid employment at Serimmune, board membership at Serimmune, and patent applications on behalf of Serimmune.

## Background

The role of adaptive immunity in cancer is of great translational interest given the recent development of novel, clinically effective immunotherapies that focus on generating T cell responses to tumor antigens. While the T-cell landscape of numerous cancer types has been explored in some depth, the role of humoral immunity in cancer is much less well-characterized. Several studies have demonstrated that a distinct antibody signature may be detectable in the serum of breast^1^, prostate^2^, and lung^3^ cancer patients and may thus be useful for cancer detection. Additionally, studies have demonstrated that B-cell infiltration into the tumor microenvironment is associated with prolonged patient survival and enhanced response to immunotherapy in melanomas, renal cell carcinomas, and sarcomas^4–9^, with several studies suggesting that B-cell autoantibodies may play a direct role in mounting an anti-tumor response^10,11^. In the setting of cancer vaccines, preclinical data indicate that IgG anti-tumor antibody responses to neoantigens in a mouse model of breast cancer can predict corresponding T cell responses to the same epitopes^12^. Furthermore, in a completed phase III trial that led to approval of the autologous cellular vaccine sipuleucel-T for mCRPC, which was one of the first immunotherapies approved by the FDA for solid tumors, productive antibody responses to the immunogen were correlated with longer overall survival in retrospective analysis^13^. Finally, anti-tumor immune responses can also be stimulated by proteins ectopically expressed outside of immune-privileged sites in somatic tumor tissues, the prototype of which is cancer-testis antigen NY-ESO-1. The prevalence of autoantibodies to the NY-ESO-1 peptide and putative conservation of B- and T-cell epitopes has led to over 30 NY-ESO-1 T-cell receptor immunotherapy clinical trials, at various stages of completion, in diverse cancer types^14,15^. Altogether, these findings support the notion that a patient’s antibody repertoire may reflect a specific immune response to the patient’s cancer and may have potential diagnostic and therapeutic implications.

Tumor-associated antibodies detectable in patient serum are traditionally profiled using microarray-based methods^16–18^, phage-display approaches^19–21^, or techniques incorporating principles of the two^22–24^. One key limitation of candidate protein-based approaches is the throughput and subsequently limited number of antigens that can be profiled and the inability to detect patient- or tumor-specific sequence variants generated by mutation. The serum epitope repertoire analysis (SERA) tool leverages a randomized bacterial-display library paired with next generation sequencing (NGS) to identify peptides binding to serum antibodies^25^. By leveraging the randomized library, SERA is able to examine both wild type and mutant sequences without any modification to the experimental protocols. Protein-based Immunome Wide Association Study (PIWAS) builds on top of the SERA assay to identify proteome-constrained antigenic signals from the SERA assay. PIWAS calculates, for each sample and protein, a smoothed log-enrichment value across a window of overlapping kmers. By comparing PIWAS values between cohorts using the outlier sum, PIWAS has been shown to identify autoantigens against the human proteome^26^.

While autoantibody enrichment has previously been demonstrated in prostate cancer, these studies were limited by smaller discovery cohorts^27,28^ or relatively restrictive peptide libraries^29,30^. It also appears that autoantibody enrichment may be context-specific. For example, one large study that leveraged a phage-display approach developed a signature for prostate cancer screening but found that this signature could be found only in a minority of patients with castration-resistant disease^2^. Thus, the autoantibody landscape for patients with metastatic castration-resistant prostate cancer (mCRPC) has yet to be elucidated.

Given that metastatic castration-resistant prostate cancer (mCRPC) represents one of the leading causes of cancer-associated death in men, we sought to characterize the autoantibody landscape of this disease. Utilizing SERA, PIWAS, and IMUNE^25,31^, we performed an unbiased analysis of autoantibodies enriched in the serum of mCRPC patients compared to healthy controls.

Specifically, we leveraged DNA-sequence level information from the assay to identify not only the proteins but also the specific epitopes (sub-peptides) within the full-length proteins that were putatively antigenic in mCRPC. We also integrated the serum antibody-profiling results with whole-genome sequencing performed on metastatic tumor biopsies and peripheral blood (germline) specimens from the same patients to assess the immunogenicity of antigens resulting from somatic mutations. We validated our top candidate antigen in *NY-ESO-1*, a known immunogenic tumor marker across cancer types, using an independent cohort of melanoma patients. We further validated the PIWAS-based seropositive results of our top motifs using ELISA experiments performed on the same serum specimens. In total, our study both recovered previously identified cancer antigens and identified novel, putative cancer-specific antigens in mCRPC.

## Materials and Methods

### Data acquisition and sample processing

A prospective IRB-approved study (NCT02432001) was conducted by a multi-institutional consortium that obtained serum, peripheral blood, and fresh-frozen, image-guided biopsy samples of metastases from mCRPC patients. Serum samples for each patient were prospectively obtained at time of study enrollment, at three-month follow-up, and at time(s) of cancer progression, if applicable. Blood was drawn at start of therapy, 3 months into therapy, and at clinically determined disease progression in serum separator tubes of 6mL (BD #367815) or 10mL (BD #367820). Tubes were spun within 90 minutes of collection (1500rcf for 10min), aliquoted into 2mL cryovials, and frozen on dry ice and shipped to a central lab at UCSF. Vials were stored upon arrival at −80°C until batch shipping on dry ice for processing at Serimmune.

Solid-tumor metastases biopsies were sequenced using whole-genome sequencing and RNA-seq as previously described^32,33^. Serum samples from a control group consisting of 1,157 individuals without known history of cancer or other predicate disease were obtained from the Serimmune database of samples. A cohort of 106 melanoma patients was used for validation of specific antigens. The prospective IRB-approved study (11-003254) of these patients was conducted at University of California Los Angeles (UCLA) that obtained peripheral blood and biopsy samples for various analysis from patients treated for advanced melanoma malignancies. Plasma samples for each patient were prospectively obtained at time of study enrollment, at approximately three-month follow-up and at further follow-up time(s) as prescribed. At baseline and after approximately 3 months of treatment, blood was collected in K3-EDTA lavender tubes of 9mL (Greiner Bio-One# 455036) and so forth. Tubes were spun within 24-hours after collection (1200rcf for 10 min, brake off), aliquoted at 500uL into 2mL cryovials for long term storage at - 80°C. A total of 106-subject aliquots were prepared as 120uL and overnight shipped on dry ice for processing at Serimmune.

### Serum antibody-epitope profiling

An *E. coli* bacterial-display library consisting of plasmids encoding random 12-mer peptides at a diversity of 8×10^9^ was constructed and prepared as previously described^25^. Serum samples were screened on this library as previously described^26^. Briefly, serum samples, at a 1:25 dilution, were added to each well of a 96 well deep well plate containing 8×10^10^ (10-fold over-sampling) induced library cells and incubated with orbital shaking at 4°C for 1 hour. Cells were washed once with PBS containing 0.05% Tween-20 (PBST) and then incubated with Protein A/G Sera-Mag SpeedBeads (GE Life Sciences, 17152104010350) for 1 hour at 4°C with orbital shaking. Cells displaying peptides bound to serum IgG antibodies were captured by magnetic separation and washed five times with PBST. Selected cells were grown overnight in LB supplemented with 34 μg/mL chloramphenicol and 0.2% wt/vol glucose at 37°C with shaking at 250 rpm.

Amplicon preparation and NGS sequencing were performed as previously described^26^. Briefly, plasmids were isolated from selected library cells using the Montage Plasmid MiniprepHTS Kit (MilliPore, LSKP09604) on a MultiscreenHTS Vacuum Manifold (MilliPore, MSVMHTS00) following the manufacturer’s instructions. Next, DNA encoding the 12mer variable regions was amplified and barcoded by two rounds of PCR. Finally, after normalizing DNA concentrations, pooled samples were sequenced using a NextSeq 500 (Illumina) and a High Output v2, 75 cycle kit (Illumina, FC-404-2005) with PhiX Run Control (Illumina, FC-110-3001) at 40% of the final pool concentration.

### Identifying mutation-specific epitopes

Previously-published results of whole-genome sequencing performed on fresh-frozen metastasis biopsies and paired peripheral blood samples of the same patients^32,34^ was analyzed to identify somatic protein-coding point and frameshift mutations present in each patient’s tumor. Data from the SERA platform were broken into 5mers and 6mers for every sample and enrichments were calculated^25^. Using the same approach as PIWAS, these enrichments were tiled against both the wild type and mutated protein sequences^26^. The enrichment values for the wild type sequence were subtracted from the mutant sequence to identify differential signal. The maximum differential value was calculated for every mutated protein and the associated patient sample. The data were fit to an exponential distribution and the probability density function was used to estimate *P*-values for every protein. *P*-values were corrected for multiple hypothesis testing using the Benjamini-Hochberg procedure^35^.

### PIWAS approach

Using the prostate cancer patients as cases and the individuals without known cancer as controls, we ran a PIWAS analysis against the human proteome^26^. PIWAS was parameterized to have a window size of 5, the number of standard deviation approach, and the maximum peak signal. The outlier sum false discovery rate as defined previously was used to prioritize antigens^26^. The reference human proteome was downloaded from Uniprot on February 28, 2019. The validation PIWAS was run using the same parameterizations with the melanoma cohort as cases and the individuals without known cancers as controls.

### Panel score approach

For top antigens *NY-ESO-1* and *HERV-K*, additional steps were taken to develop a motif panel for these antigens. In both cases, the following approach was taken. First, prostate cancer samples with an antigen PIWAS score >6 were identified. The positive prostate cancer samples and 30 random healthy controls were used as input to the IMUNE algorithm, which was parameterized with 20% sensitivity and 100% specificity^25^. Motifs that mapped linearly to the target antigen were retained. For each retained motif, the mean and standard deviation (SD) of enrichment scores was calculated using the 1,157 control specimens as a reference group. z-scores were calculated for every cancer specimen. Then, for each cancer specimen, enrichment scores for each motif were z-scored (based on the enrichment score mean and SD of the control group) and summed to generate a composite score for each specimen. Thus, the final composite “panel score” was defined as the sum of motif z-scores for each specimen. Thresholds for positivity on the panel were set at a 99% specificity.

### ELISA

Briefly, *NY-ESO-1* recombinant protein (Origene) at 0.5 ug/ml, or control recombinant protein CENPA (Origene) at 0.5 ug/ml, or *HERVK-5* recombinant protein (MyBiosource) at 1 ug/ml or control protein Bovine Serum Albumin (Sigma) at 1 ug/ml in PBS were coated onto flat bottom, 96 well plates (Nunc MaxiSorp), 50 ul per well at 4°C overnight. Plates were washed with PBS containing 0.1% Tween 20 and blocked with 5% non-fat milk in PBS for 2 hours at room temperature. Plates were then incubated with 100 ul of patient serum diluted 1/200 or 1/2000 in 5% non-fat milk in PBS for 2 hours at room temperature. Following washing, plates were incubated with peroxidase conjugated goat anti-human IgG secondary (1/10,000 in 5% non-fat milk in PBS; Jackson ImmunoResearch) for 1 hour at room temperature. After a last wash step, the reaction was developed with 3,3′,5,5′-tetramethylbenzidine substrate solution (ThermoFisher) for 1-10 minutes and stopped with 1M hydrochloric acid. Absorbance at 450 nm was measured on a plate reader. ELISA values were calculated as the mean difference between the testing recombinant protein and the control protein. Due to reagent availability constraints, sera reactivity to *HERVK-5* was assessed in lieu of reactivity of *HERVK-113* given the high sequence similarity of the *HERVK-5* and *HERVK-113* proteins (95.2% per BLAST analysis).

### Statistical methods and survival analysis

Overall survival was measured from time of mCRPC diagnosis. Survival analyses were conducted using the Kaplan-Meier method with log-rank testing for significance. The χ^2^ test was performed to assess the relationship between ELISA and PIWAS antibody enrichment results. All independence and hypothesis tests were performed using a two-sided significance level of 0.05. Multiple hypothesis testing correction was performed using the Benjamini-Hochberg procedure.

## Results

Serum specimens were obtained from a cohort of 72 mCRPC patients with a mean age of 72 years at time of mCRPC diagnosis (**Table 1**). The cohort was predominantly Caucasian (87%) with high-grade primary tumors in 54%.Visceral metastases were observed in 15 of 72 (21%) patients. Sera obtained at more than one timepoint were available for 79% of patients (**Table S1)**.

**Table 1:**
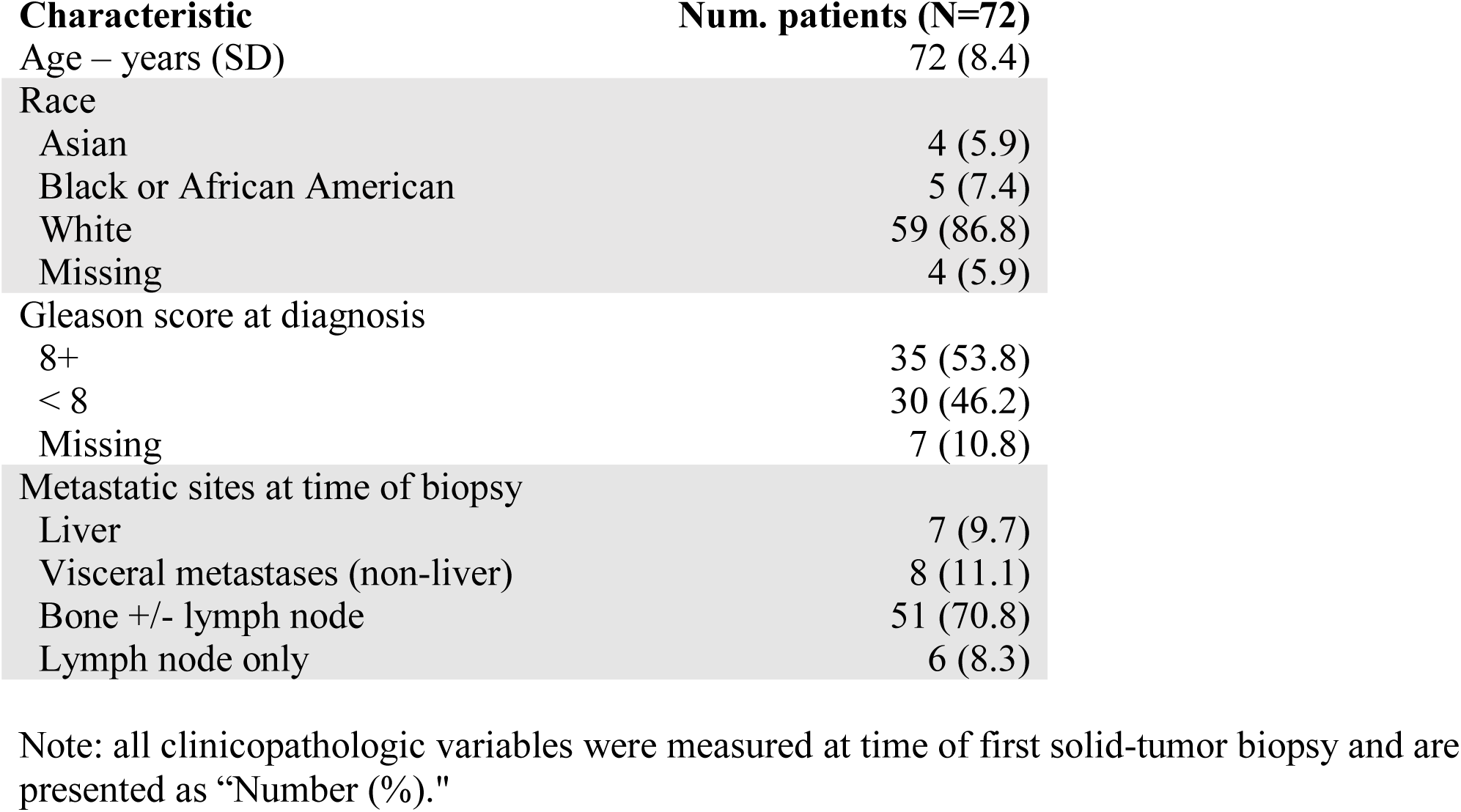
Patient clinicopathologic characteristics.

| **Characteristic** | **Num. patients (N=72)** |
| --- | --- |
| Age – years (SD) | 72 (8.4) |
| Race |  |
| Asian | 4 (5.9) |
| Black or African American | 5 (7.4) |
| White | 59 (86.8) |
| Missing | 4 (5.9) |
| Gleason score at diagnosis |  |
| 8+ | 35 (53.8) |
| < 8 | 30 (46.2) |
| Missing | 7 (10.8) |
| Metastatic sites at time of biopsy |  |
| Liver | 7 (9.7) |
| Visceral metastases (non-liver) | 8 (11.1) |
| Bone +/- lymph node | 51 (70.8) |
| Lymph node only | 6 (8.3) |

Note: all clinicopathologic variables were measured at time of first solid-tumor biopsy and are presented as “Number (%).”

Integrating serum antibody profiling results with whole-genome sequencing results, we first sought to assess whether somatic, protein-coding mutations were associated with an antibody response specific to the mutant peptide in mCRPC. 29 of the 6,636 protein-coding somatic mutations observed in our cohort were associated with a significant enrichment (exponential FDR < 0.05) in antibodies specific to the mutated peptide (**Figure 1A, Table S1**). These 29 mutations were approximately evenly distributed between frameshift and missense mutations (**Figure S1A**). These events constituted the minority of mutations (0.44%), consistent with literature that suggests that most protein-coding mutations do not elicit an immune response^36^. Each of the mutation-specific antibodies was enriched in only one patient. However, the somatic mutations that coded the epitopes were also private to individual patients. This suggested that the observed antibody response was specific to the individual in which the mutant antigen was available. In 11 of 20 mutant epitopes derived from patients with multiple serum specimens available, multiple independent serum samples obtained from the same individual at different timepoints confirmed the mutation-specific antibody enrichments (**Table S2**). We highlight one example of a patient with a point mutation and a patient with a frameshift mutation that demonstrated an enriched autoantibody response to the corresponding mutant epitope across multiple timepoints (**Figures 1B, 1C**).

**Figure 1.**
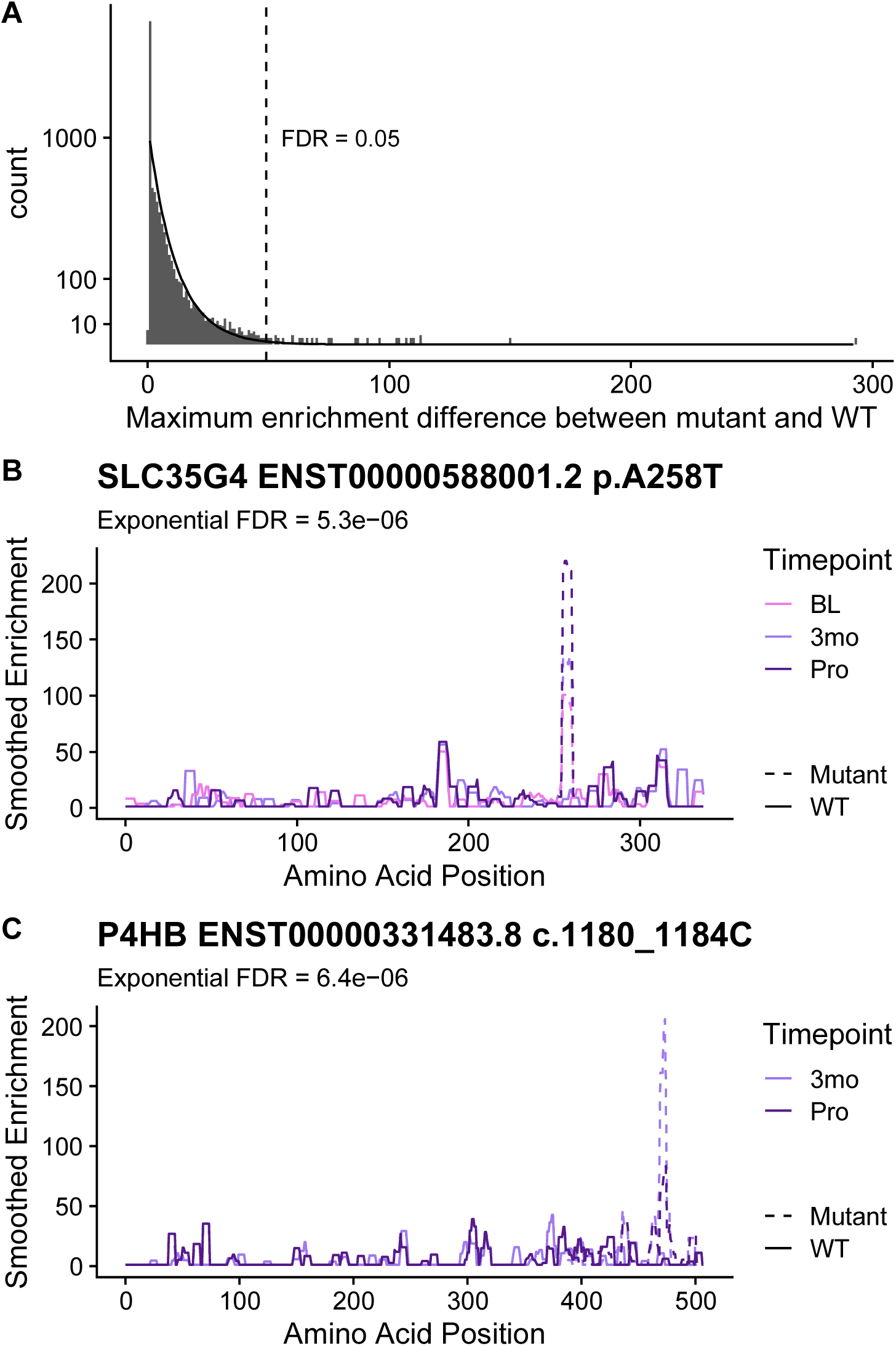
Analysis of mutation-specific epitopes in the prostate cancer patients. (A) Enrichments were calculated for every mutated protein in the affected patients (n mutations = 6,636; n patients = 72). An exponential distribution (indicated by the line) was fit to the data to calculate significance of each mutation. Difference between enrichment values in the mutant sequence and the wild type sequence in that patient are shown along the x-axis. Distributions based on mutation type are shown in **Figure S1**. (B) A high scoring point mutation in SLC35G4 is shown for patient DTB-129. (C) A high scoring frameshift mutation in P4HB is shown for patient DTB-102. (BL, baseline; 3mo, 3months after enrollment in study; Pro, disease progression)

Next, to investigate cancer-specific autoantibodies resulting from non-mutant proteins, we performed a protein-based immunome-wide association study (PIWAS). We found 11 proteins to be significantly enriched for antibodies in mCRPC patients compared to healthy controls (**Figure 2, Table 2/S3**). The top two candidates were cancer-testis antigens *NY-ESO-1* and *NY-ESO-2*, with the dominant epitope occurring in a conserved region between the proteins. Eight of 72 (11%) patients demonstrated PIWAS values > 6, all of which mapped to amino acids 11-30 of *NY-ESO-1* (**Figures 3A, S3A)**. PIWAS values for seven of these eight patients remained above the threshold at all timepoints (**Figure S3B**). Of note, this dominant B-cell epitope had been described in a previous study using a peptide approach and was found to be present in prostate cancer at a similar frequency^37^. In order to identify additional *NY-ESO-1* antigenic regions, we applied the previously described IMUNE algorithm to identify peptide motifs that were significantly enriched in prostate cancer patients relative to healthy controls^25^. For this analysis, eight *NY-ESO-1* PIWAS positive samples were analyzed by IMUNE using 30 healthy patient samples as controls. A total of nine cancer-specific motifs were identified that mapped to *NY-ESO-1* (**Figure 3B**). While seven of the nine motifs aligned to the same portion of *NY-ESO-1* identified by PIWAS, two of the motifs align to a new epitope around that 100^th^ amino acid that is additionally present in samples without the PIWAS epitope. Samples with composite panel scores greater than 6.6 (based on a pre-defined 99% specificity threshold) were designated positive. Using this panel, nine of 72 patients (12.5%) including one patient without enrichment of the dominant epitope were positive for *NY-ESO-1* at a specificity of 99% (**Figures 3C, S3C**).

**Table 2.**
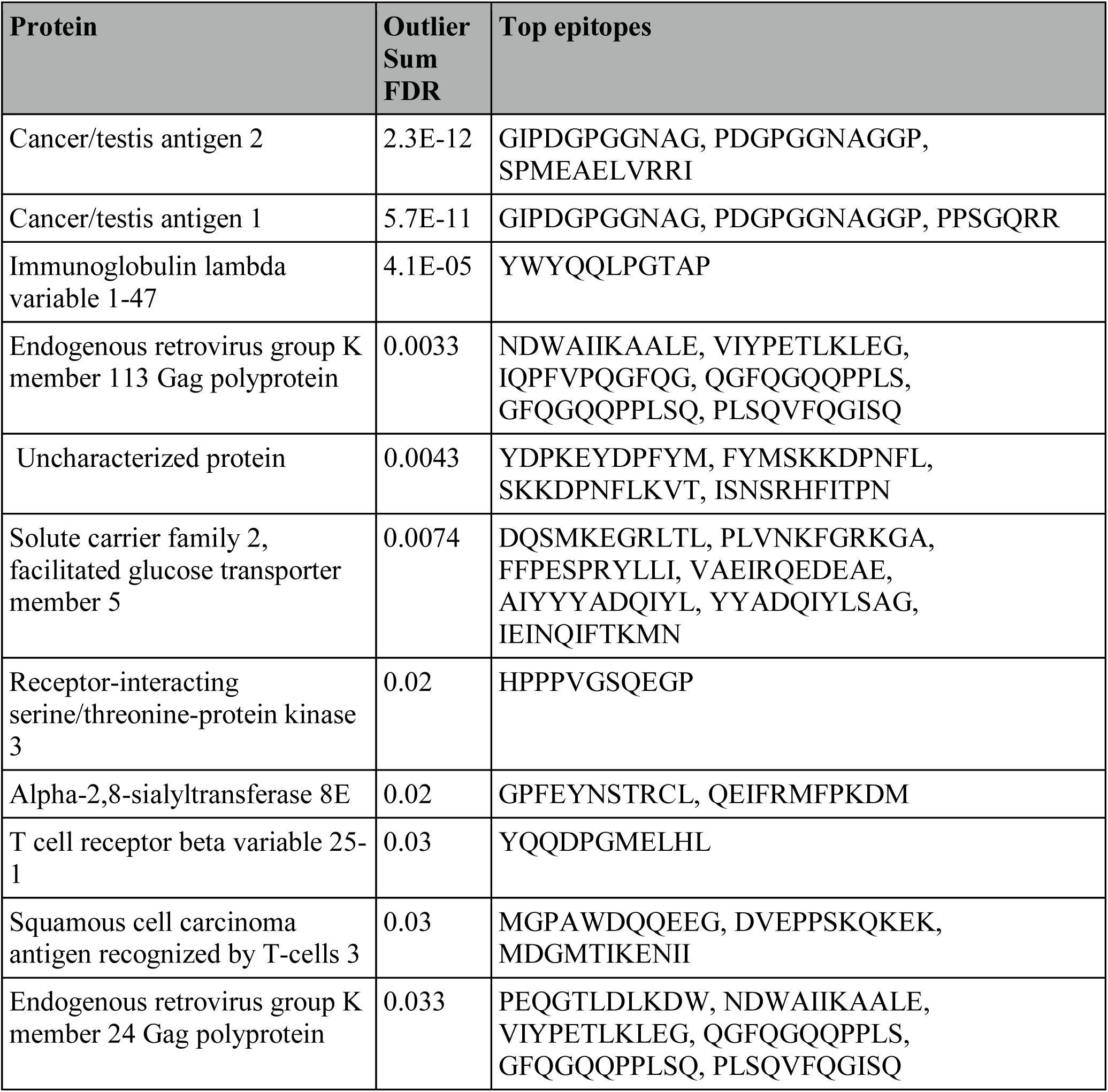
Top epitopes from prostate cancer PIWAS.

**Figure 2.**
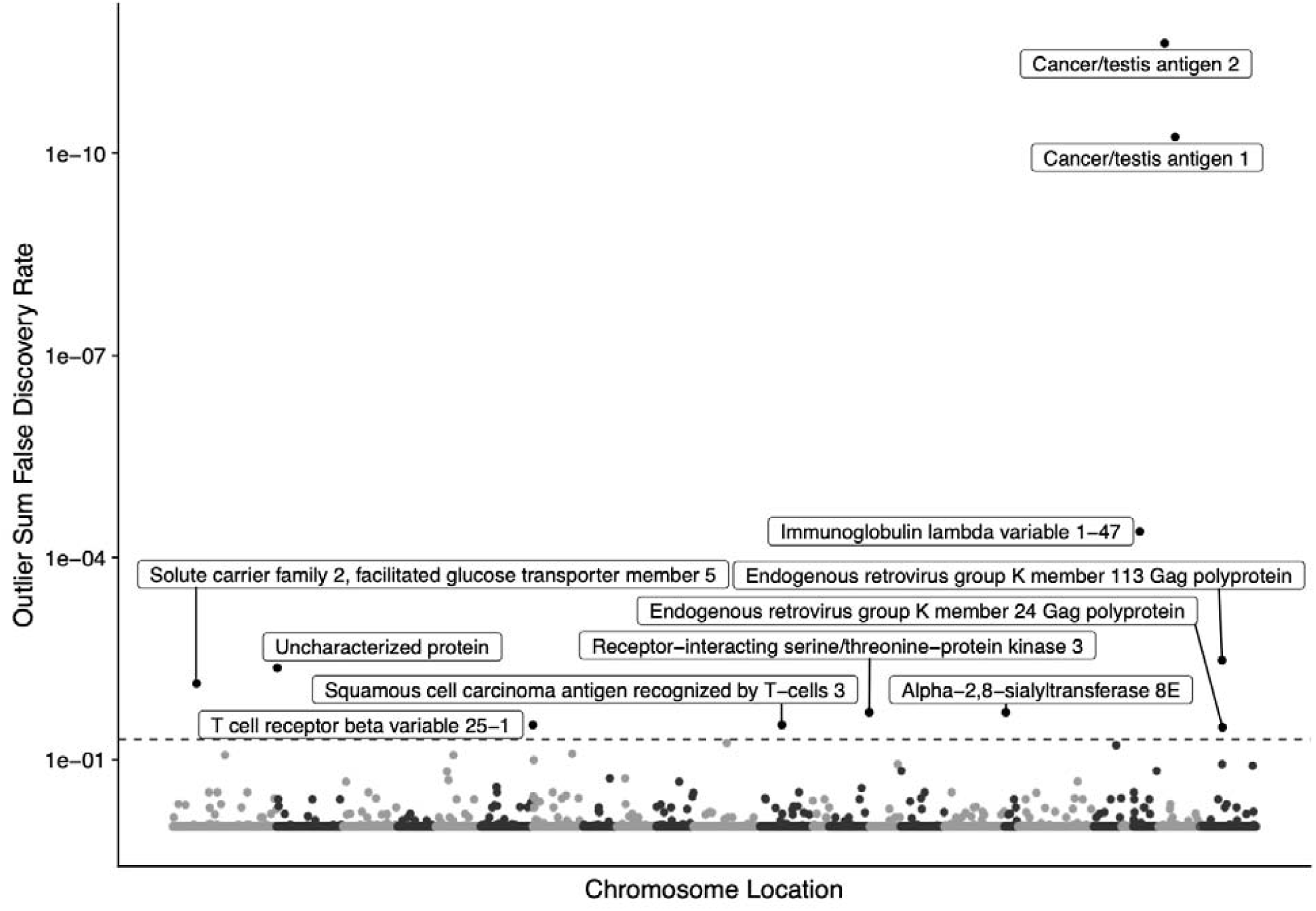
Manhattan plot of protein-based immunome wide association study (PIWAS) results highlighting antigens significantly enriched in prostate cancer compared to healthy control patients. Outlier sum FDRs are shown for every protein in the human proteome. Labels are shown for all proteins with an FDR < 0.05. Co-positivity of proteins with FDR < 0.05 are shown in Figure S2.

**Figure 3.**
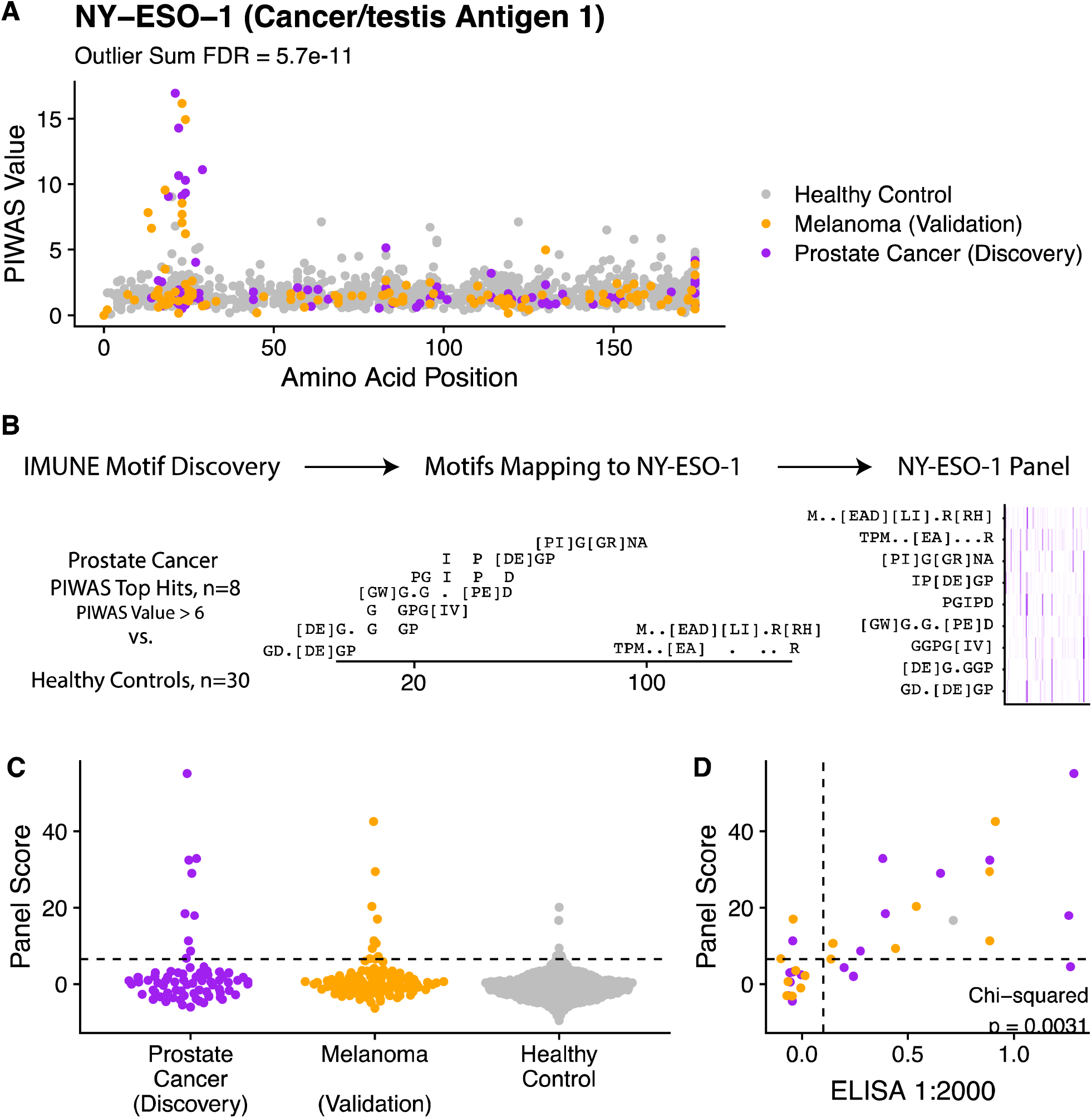
Discovery and validation of *NY-ESO-1* antigenic signal. (A) Manhattan plot visualizing PIWAS values for *NY-ESO-1*, with one point per patient being shown and colored by patient subgroup (purple = prostate cancer (discovery, n=72), orange = melanoma (validation, n =218), (grey = controls without known disease (n=1,157)). (B) Prostate cancer samples that are positive by PIWAS are compared to healthy controls using IMUNE motif discovery algorithm. Motifs which map linearly to NY-ESO-1 are retained. A panel score is calculated by summing enrichment z-scores across all motifs. (C) Dot plot of panel score for serum specimens stratified by patient subgroup. (D) Scatterplot demonstrating concordance between *NY-ESO-1* panel score and ELISA results, assessed using the Chi-square test of independence. Points are colored based on patient subgroup. Additional results provided in Figure S3.

To validate this finding with an orthogonal serum profiling approach, the composite panel score results were benchmarked against a NY-ESO-1 ELISA experiment performed on the same prostate cancer serum samples. We found that the panel score and ELISA results were strongly associated (Cohen’s kappa = 0.57, **Figure 3D**). RNA-seq expression data revealed that NY-ESO-1 was expressed in the metastases of six of nine patients demonstrating *NY-ESO-1* antibody enrichment at time of initial metastatic tumor biopsy, confirming antigenic availability in these patients. Altogther, these findings demonstrated the high sensitivity and specificity of the joint PIWAS-IMUNE (PIWAS-I) approach in identifying disease-specific epitopes in prostate cancer. While cancer-specific, the dominant *NY-ESO-1* epitope was previously known to be a tumor marker in not only advanced prostate cancer but also other cancers including melanoma. Specifically, a prior study found autoantibodies to the dominant *NY-ESO-1* epitope to be enriched in 12.5% of melanoma samples^37^. To assess whether a similar finding would be observed using the PIWAS-I approach, we profiled the serum antibody repertoires of an independent cohort consisting of 106 melanoma patients. We observed the dominant *NY-ESO-1* epitope enriched in 8 of 106 (7.5%) samples (**Figures 3A, 3C**), consistent with the prior report. This finding further validated the robustness of the PIWAS-I approach.

In addition to NY-ESO-1, the HERV (HML-2) family of proteins were also found by PIWAS to be significantly enriched for autoantibodies in mCRPC. 9 of 72 (12.5%) patients demonstrated significant enrichment of autoantibodies to HERVK-113, with recurrent epitopes identified near the 155^th^ amino acid and C-terminus of the protein (**Figures 4A, S4A**). The presence of autoantibody enrichment to HERVK-113 was consistent across different timepoints in patients with multiple independently sampled serum specimens (**Figure S4B**). A motif panel for HERVK-113 was generated as described above using 9 PIWAS positive samples for IMUNE-based motif discovery (**Figure 4B**). The panel scores were enriched in 16 of 72 (22.2%) patients (**Figures 4C, S4C**). The panel scores were highly concordant with confirmatory ELISA testing (Cohen’s kappa = 0.91, **Figure 4D**). Additionally, RNA-seq expression of HERVK-113 was observed in all patients panel-score positive for HERVK-113, suggesting antigenic availability.

**Figure 4.**
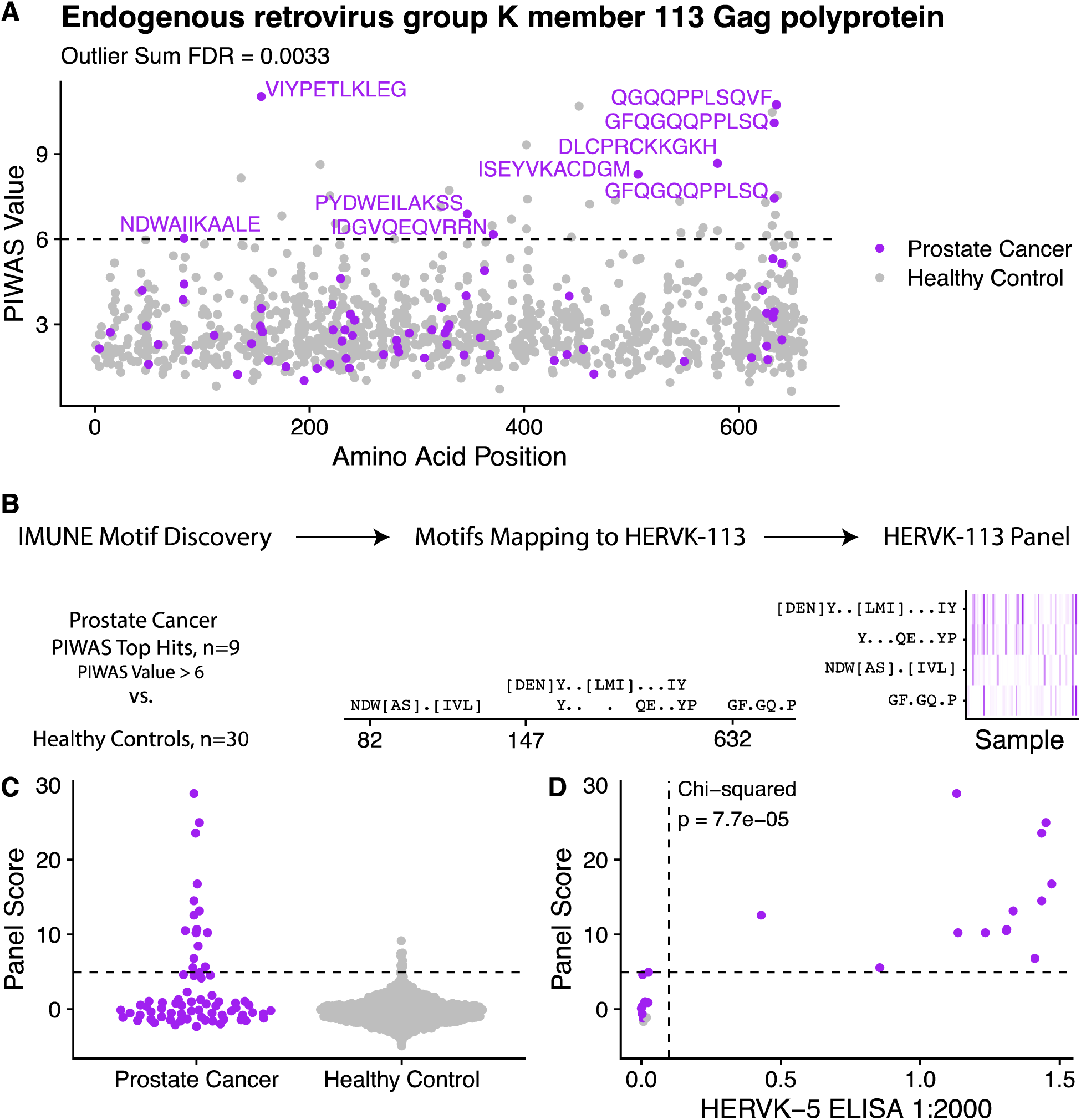
Discovery and validation of *HERVK-113* antigenic signal. (A) Manhattan plot visualizing antibody-enrichment scores for *HERVK-113*. Epitopes associated with samples with a PIWAS value greater than 6 are labeled. (B) Prostate cancer samples that are positive by PIWAS are compared to healthy controls using IMUNE motif discovery algorithm. Motifs which map linearly to HERVK-113 are retained. A panel score is calculated by summing enrichment z-scores across all motifs. (C) Dot plot of panel scores for prostate cancer patients compared to healthy controls. (D) Scatterplot of HERVK-113 panel score vs. HERVK-5 ELISA titer score. Additional results provided in Figure S4.

Additional mCRPC-specific epitopes were identified using the PIWAS approach. These included epitopes in the protein products of *SART3, RIPK3, ST8SIA5, IGLV1-47, TRBV25-1*, and SLC2A5 (**Figure 2, Table S3**). Seven of 72 (9.7%) patients demonstrated significant enrichment of autoantibodies to *SART3*, 6 of 72 (8.3%) demonstrated enrichment of autoantibodies to *RIPK3*, 5 of 72 (6.9%) demonstrated enrichment of autoantibodies to *ST8SIA5*, and 4 of 72 (5.6%) demonstrated enrichment of autoantibodies to *IGLV1-47* (**Figure S2**). To assess autoantibody co-enrichment patterns, we assessed pairwise correlation between autoantibody enrichment scores of the eleven putative mCRPC-specific epitopes using our cohort of 72 patients (**Figure S2A-B**). Enrichment of autoantibodies to NY-ESO-1 and HERVK-113 was mutually exclusive on PIWAS analysis, as no patients demonstrated significant antibody enrichment to both NY-ESO-1 and HERVK-113 (**Figure S2A**). Presence of autoantibody enrichment to epitopes in NY-ESO-1 or HERVK-113 were not prognostic of overall survival (**Figures S3D, S4D**).

## Discussion

Herein, we have characterized the autoantibody landscape of metastatic castration-resistant prostate cancer. We observed cancer-specific enrichment of antibodies to mutant peptides in select genes and to non-mutant peptides in the *NY-ESO-1* and *HERVK-113* proteins among others.

Previous reports demonstrated that disease-specific neoepitopes may include defective gene products resulting from somatic alterations such as mutations^38^ and errors in protein translation^39^. The extension of this principle to cancer is supported by prior studies in lung and colorectal cancers, which found that tumors with missense mutations in *TP53* and frameshift mutations in select genes were associated with autoantibodies to the mutant protein products^40,41^. We performed the first comprehensive assessment of cancer-specific B cell neoantigens to date and observed several examples of this phenomenon in genes such as *SLC35G4* and *P4HB*. However, the majority (99.6%) of somatic mutations did not result in antibody-specific epitopes in our cohort. This finding is consistent with prior T-cell studies, which found that only a minority of mutations stimulate a specific T-cell response^36,42^. In the present study, mutations generating the strongest detected responses were approximately equally distributed between missense and frameshift mutations.

The mutations that were observed to be associated with an epitope-specific humoral immune response tended to be private to individual patients rather than shared among multiple individuals. This observation too is consistent with a prior report in colorectal cancer^42^ and may be due to the fact that somatic mutations themselves (and hence the resulting aberrant mutant protein) tend not to be recurrent across mCRPC patients. Nevertheless, we observed that the autoantibodies to mutant peptides were often present in multiple serum specimens collected independently from the same patient. These data validate the specificity of the epitope profiling and PIWAS approaches and support the notion that select mutations may induce a humoral immune response in mCRPC.

We observed enrichment of autoantibodies to not only mutant but also wild-type epitopes. This is supported by prior studies which suggest that cancer-specific overexpression of non-mutant antigens may comprise the majority of tumor-associated antigens^43–46^. *NY-ESO-1*, or cancer testis antigen 1B (*CTGAG1B*), has been well-characterized as an antigen that elicits humoral immune responses in various cancers including melanoma and breast, lung, bladder, ovarian, and prostate cancers^47–49^. Additionally, given its cancer-specific expression pattern outside of the testes^49–52^, *NY-ESO-1* has shown great promise as a potential target for T-cell immunotherapies in various cancers^47^. In our unbiased approach to identifying immunogenic antigens, we found that *NY-ESO-1* was the top candidate. Moreover, by experimental design, we were able to identify the specific recurrent motif that has been previously demonstrated to be immunogenic in multiple cancer types^37,53^. By leveraging the PIWAS-I approach, we were additionally able to identify motifs which do not contain continuous matches to the protein sequence, improving both sensitivity and specificity. These findings, along with empiric validation via the ELISA approach, support the notion that PIWAS-I can be used to reliably recover immunogenic motifs in cancer.

The PIWAS-I approach also identified epitopes in *HERVK-113*. Human endogenous retroviruses (HERVs) comprise a family of retroviruses whose genetic material has previously been integrated into the human germline and whose gene products have been implicated in cancer pathogenesis^54,55^. HERVs have been previously described as being transcriptionally activated and potentially antigenic in the context of cancer: prior studies of renal cancers and seminomas identified a cancer-specific IgG response to *ERVK-10*^22,56,57^. Humoral responses to HERVs have similarly been reported in melanomas^58^ and ovarian^59^, breast^60,61^, and prostate cancers.

Additionally, the gene product of HERV-K may be not only a biomarker of disease but also a therapeutic target, as a preclinical model demonstrated that monoclonal antibodies against the HERV-K env protein was associated with inhibition of tumor growth in breast cancer^63^. In prostate cancer particularly, detection of autoantibodies to the HERV-K gag protein has been shown to be enriched in advanced prostate cancer relative to early prostate cancer (21% vs. 1.4%) and associated with poor survival outcomes^64^. We observed a similar prevalence of 22% for HERV-K antibody enrichment in our cohort of advanced prostate cancer patients. However, we observed no significant difference in overall survival (OS) between mCRPC patients with and without HERV-K antibody enrichment. In contrast to the previously studied cohort, our cohort was comprised exclusively of advanced prostate cancer patients. Thus, our findings suggest that autoantibodies to the HERV-K protein may be associated with disease burden but may not be prognostic of OS amongst patients with advanced disease. Additional prospective studies are needed to explore the prognostic value of HERV-K antibody enrichment in greater detail.

Additional antigens identified through the PIWAS-I approach included *SART3* and *RIPK3.* Both of these genes have been previously implicated as biomarkers or potential regulators of cancer progression. *SART3* is a cancer testis antigen that is expressed specifically in various cancer tissues (excluding normal testis)^65,66^. *SART3* has been shown to induce both a humoral and cellular adaptive immune response in a vaccination study of patients with advanced colorectal cancer^67^. *RIPK3* is a tumor suppressor whose downregulation has been associated with tumorigenesis, immunomodulation, and poor clinical prognosis in colorectal cancer^68,69^, although its exact role in the adaptive immunity is still under investigation. While confirmatory studies are needed, the present study in conjunction with supporting studies in other cancer types nominates potential immune biomarkers and therapeutic targets in mCRPC.

In addition to recovering known and novel epitopes of interest in mCRPC, the findings of the present study highlight potentially conserved B- and T-cell cancer-specific epitopes and a combined B- and T-cell response to cancer. *NY-ESO-1* has been found to elicit a high-titer IgG humoral response as well as a cellular immune response in patients with melanoma^47,70^. *SART3*, or “Squamous cell carcinoma antigen recognized by T-cells 3,” was initially discovered as a T-cell epitope and was later found to also stimulate a correlated IgG response^65,67^. More generally, cancer vaccination studies have demonstrated how the humoral immune response to cancer-associated antigens may provide insights into targets of the endogenous cellular immune system^12,67^. Future work may further elucidate the frequency of epitopes on shared antigens among the humoral and cellular immune systems in mCRPC and the extent to which each contributes to antitumor activity.

In summary, we leveraged recently published epitope profiling techniques to characterize the autoantibody landscape of mCRPC and identify cancer-specific antigens and epitopes. By pairing patient serum profiling with whole-genome sequencing results of paired solid-tumor biopsies, we identified 29 novel epitopes to mutant peptides generated by patient-specific somatic mutations. We also identified 11 conserved protein antigens, with several supported by prior reports in other cancer cohorts. Our findings and the presented next generation sequencing-based approach to autoantibody profiling provide insight into immune biomarkers and potential therapeutic targets in advanced prostate cancer.

## Acknowledgements

This study was funded in part by the Stand Up To Cancer – Prostate Cancer Foundation Dream Team. MS was supported by the Swedish Research Council (Vetenskapsrådet) with grant number 2018-00382 and the Swedish Society of Medicine (Svenska Läkaresällskapet). A.R. was funded by the Parker Institute for Cancer Immunotherapy, the Ressler Family Fund, support from Ken and Donna Schultz, and NIH grants R35 CA197633 and P01 CA244118. FYF was funded by Prostate Cancer Foundation Challenge Awards and the following NIH grants: NIH/ NCI 1R01CA230516-01, NIH / NCI 1R01CA227025-01A1, NIH 2U10CA180868-06, NIH P50CA186786. We thank the Serimmune team for supporting the processing and analysis of samples, including: Minlu Zhang, Jack Reifert, Joel Bozekowski, Brian Martinez, Gregory Jordan, Timothy Johnston, Cameron Gable, Steve Kujawa, Elisabeth Baum-Jones. Special thanks to Elizabeth Stewart for editing this manuscript.

## Supplementary Materials

**Table S1:** Number of serum sample timepoints available for mCRPC cohort

| Timepoints | Number of patients (%) |
| --- | --- |
| 1 | 15 (21) |
| 2 | 31 (43) |
| 3 | 24 (33) |
| 4 | 2 (3) |

**Table S2:**
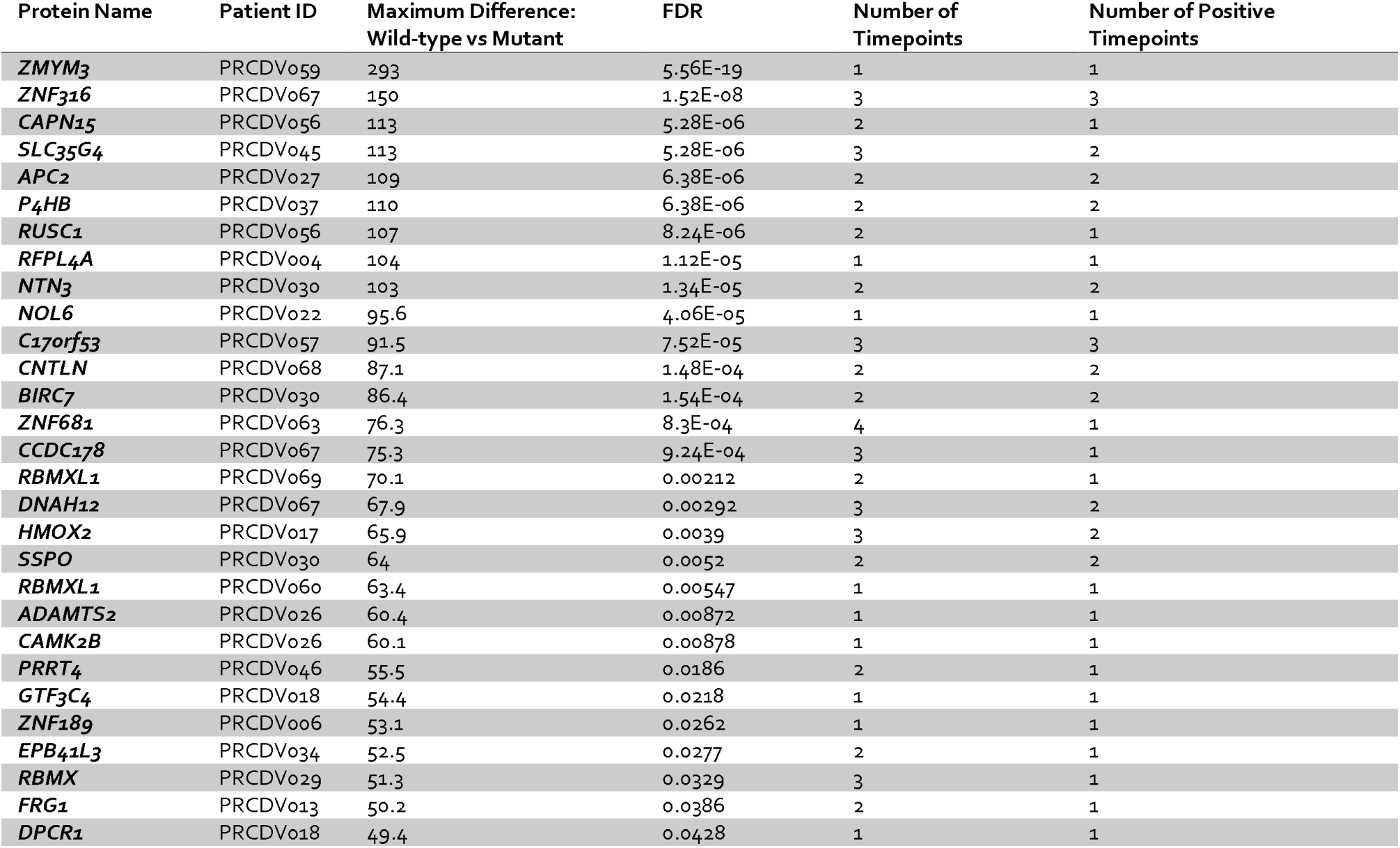
Protein and patient pairs with a significant (FDR < 0.05) difference between the wild-type and mutated sequence enrichment values. In addition, the total number of timepoints from this patient is shown alongside the number of samples from that patient that also achieve the FDR < 0.05 threshold.

| Protein Name | Patient ID | Maximum Difference:<br>Wild-type vs Mutant | FDR | Number of<br>Timepoints | Number of Positive<br>Timepoints |
| --- | --- | --- | --- | --- | --- |
| *ZMYM3* | PRCDV059 | 293 | 5.56E-19 | 1 | 1 |
| *ZNF316* | PRCDV067 | 150 | 1.52E-08 | 3 | 3 |
| *CAPN15* | PRCDV056 | 113 | 5.28E-06 | 2 | 1 |
| *SLC35G4* | PRCDV045 | 113 | 5.28E-06 | 3 | 2 |
| *APC2* | PRCDV027 | 109 | 6.38E-06 | 2 | 2 |
| *P4HB* | PRCDV037 | 110 | 6.38E-06 | 2 | 2 |
| *RUSC1* | PRCDV056 | 107 | 8.24E-06 | 2 | 1 |
| *RFPL4A* | PRCDV004 | 104 | 1.12E-05 | 1 | 1 |
| *NTN3* | PRCDV030 | 103 | 1.34E-05 | 2 | 2 |
| *NOL6* | PRCDV022 | 95.6 | 4.06E-05 | 1 | 1 |
| *C17orf53* | PRCDV057 | 91.5 | 7.52E-05 | 3 | 3 |
| *CNTLN* | PRCDV068 | 87.1 | 1.48E-04 | 2 | 2 |
| *BIRC7* | PRCDV030 | 86.4 | 1.54E-04 | 2 | 2 |
| *ZNF681* | PRCDV063 | 76.3 | 8.3E-04 | 4 | 1 |
| *CCDC178* | PRCDV067 | 75.3 | 9.24E-04 | 3 | 1 |
| *RBMXL1* | PRCDV069 | 70.1 | 0.00212 | 2 | 1 |
| *DNAH12* | PRCDV067 | 67.9 | 0.00292 | 3 | 2 |
| *HMOX2* | PRCDV017 | 65.9 | 0.0039 | 3 | 2 |
| *SSPO* | PRCDV030 | 64 | 0.0052 | 2 | 2 |
| *RBMXL1* | PRCDV060 | 63.4 | 0.00547 | 1 | 1 |
| *ADAMTS2* | PRCDV026 | 60.4 | 0.00872 | 1 | 1 |
| *CAMK2B* | PRCDV026 | 60.1 | 0.00878 | 1 | 1 |
| *PRRT4* | PRCDV046 | 55.5 | 0.0186 | 2 | 1 |
| *GTF3C4* | PRCDV018 | 54.4 | 0.0218 | 1 | 1 |
| *ZNF189* | PRCDV006 | 53.1 | 0.0262 | 1 | 1 |
| *EPB41L3* | PRCDV034 | 52.5 | 0.0277 | 2 | 1 |
| *RBMX* | PRCDV029 | 51.3 | 0.0329 | 3 | 1 |
| *FRG1* | PRCDV013 | 50.2 | 0.0386 | 2 | 1 |
| *DPCR1* | PRCDV018 | 49.4 | 0.0428 | 1 | 1 |

**Table S3:**
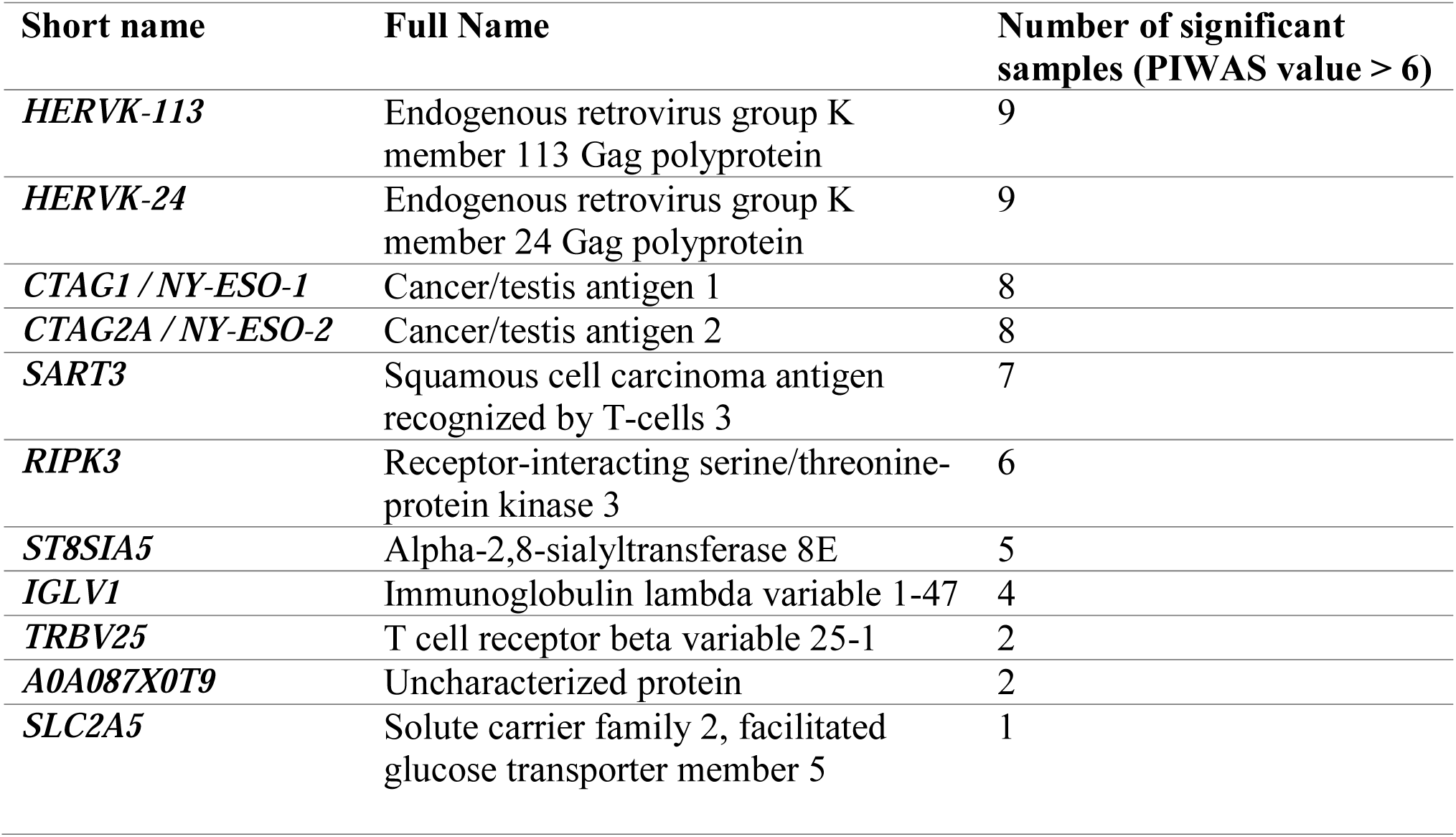
Number of PIWAS significant samples per protein in the prostate cohort

**Figure S1.**
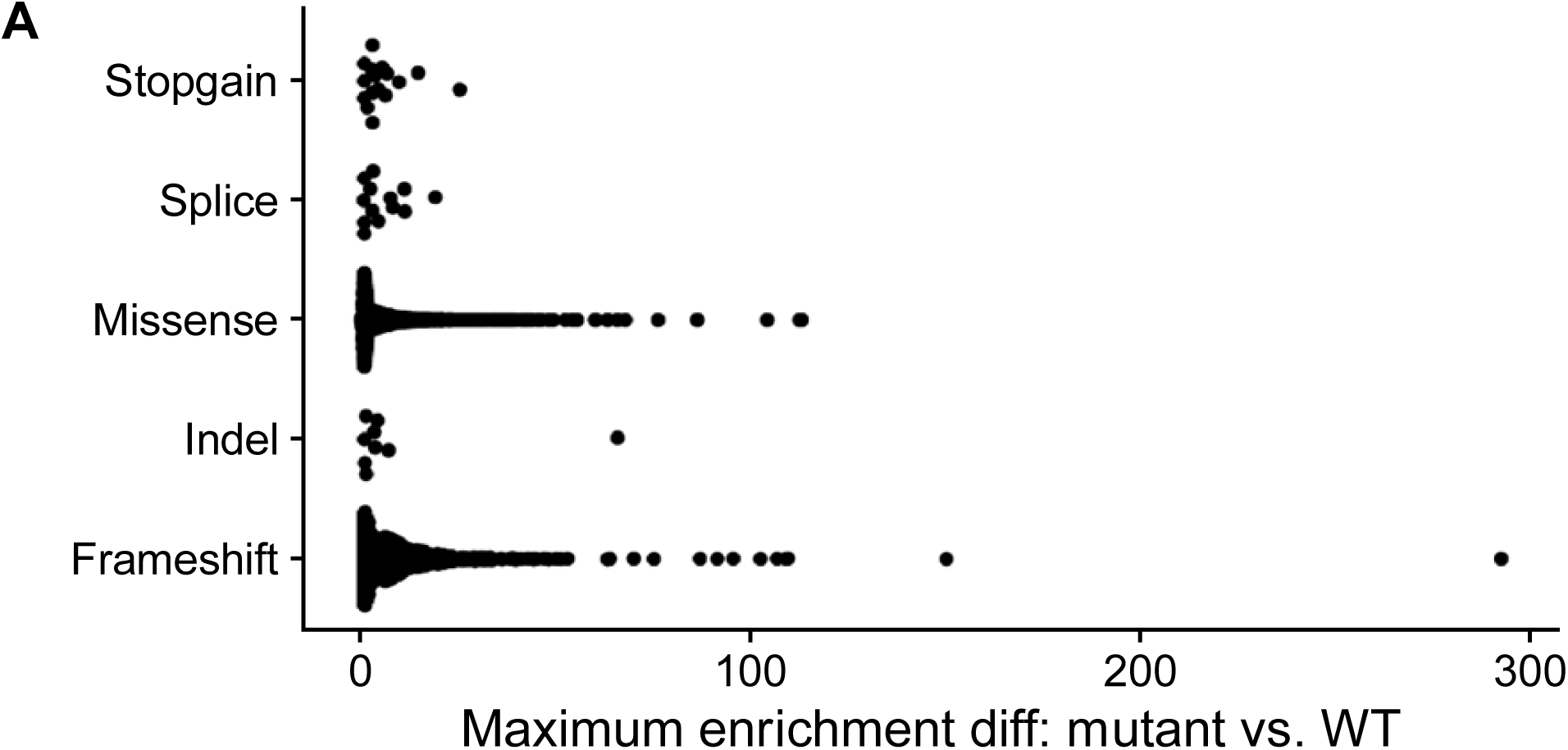
Enrichment differences in mutant vs. wild-type sequences by type of mutation.

**Figure S2.**
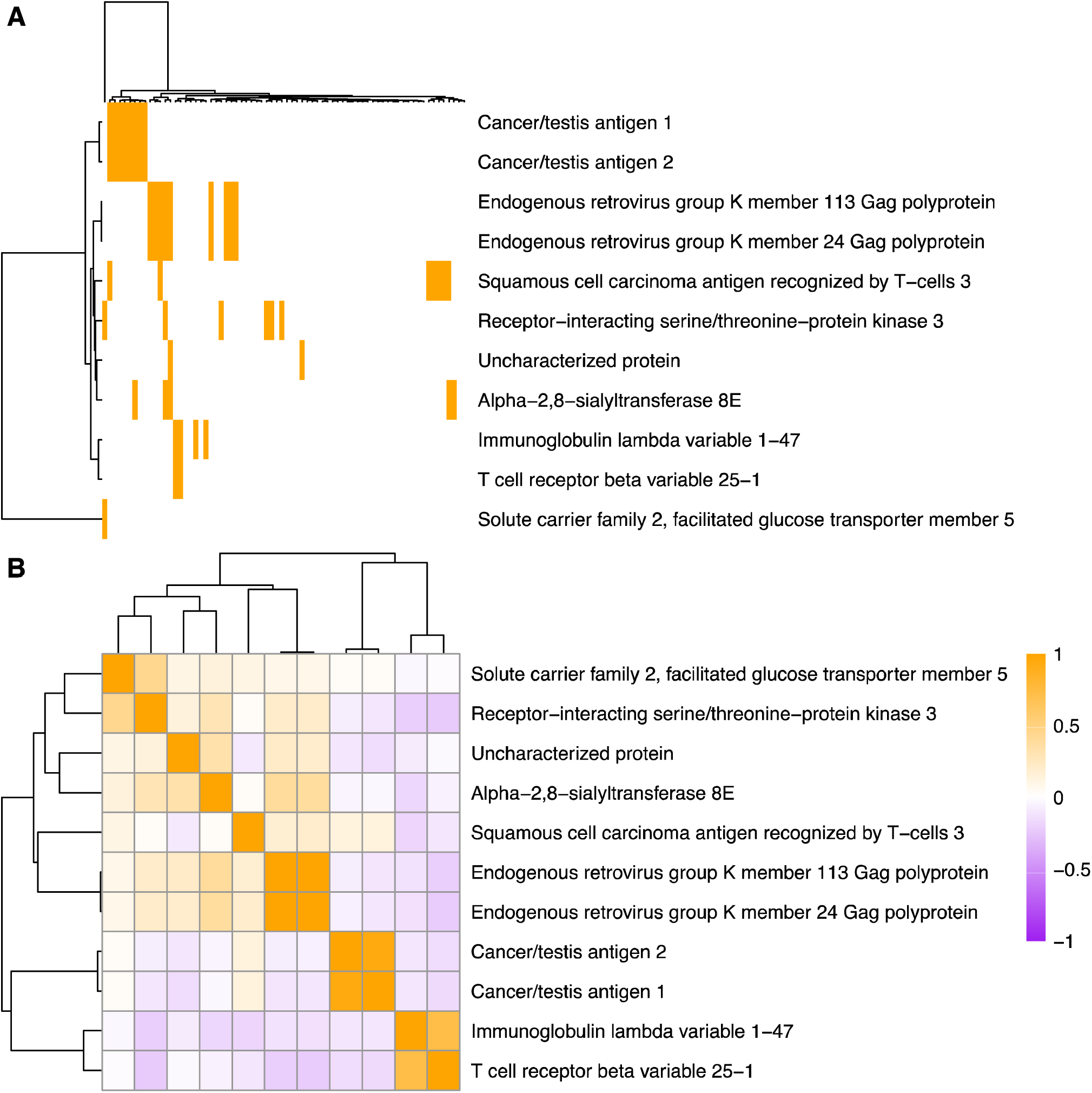
**(A)** PIWAS scores for all 72 prostate cancer patients. Orange indicates a PIWAS value greater than 6. Rows and columns are both clustered. (B) Correlations of PIWAS values across the prostate cancer samples.

**Figure S3.**
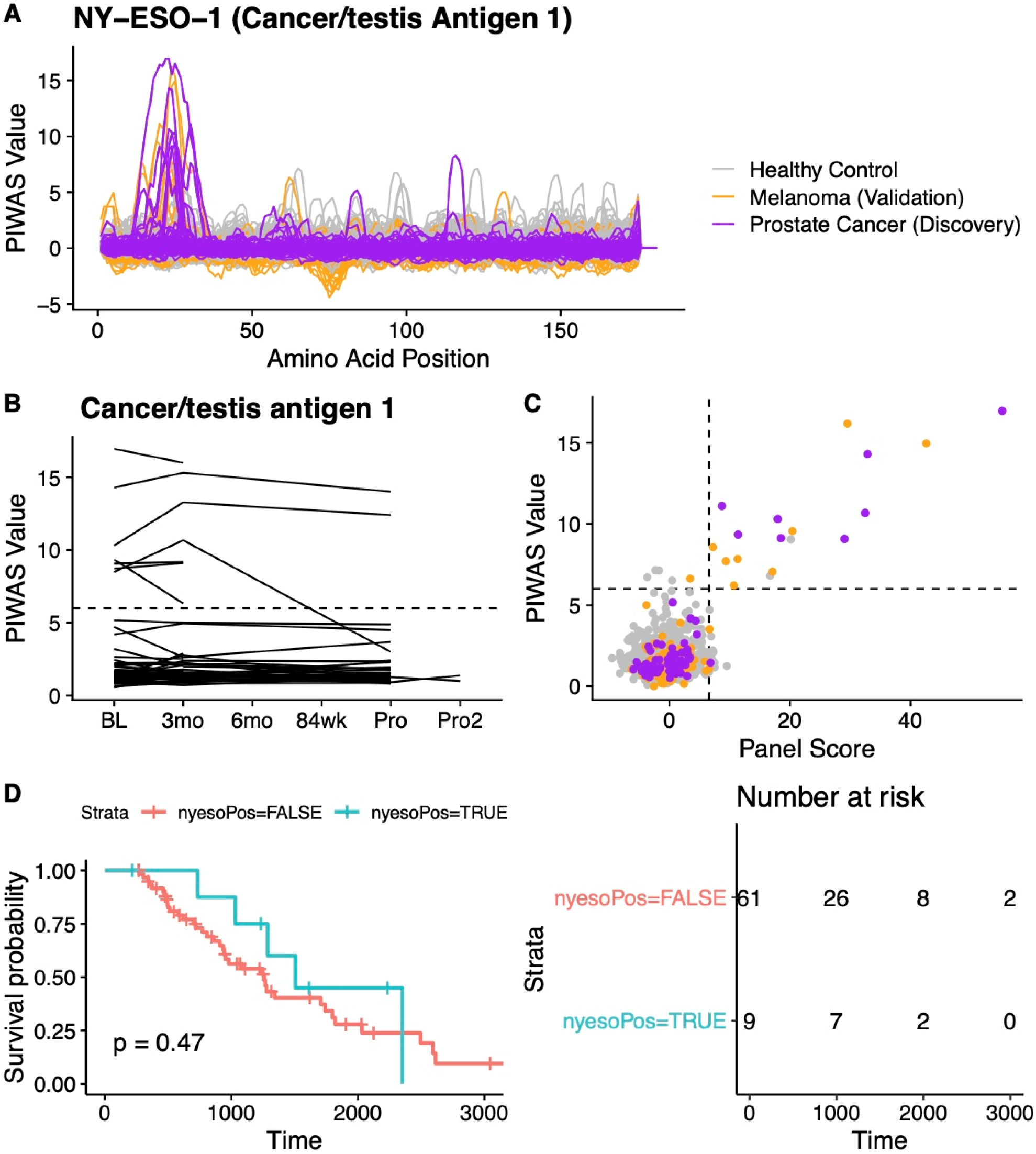
Additional findings for *NY-ESO-1*. (A) Complete tiling data for *NY-ESO-1* for every protein included in the PIWAS analysis. (B) PIWAS values across time for *NY-ESO-1*. One line is shown per patient. (C) Correlation between PIWAS value and panel score. (D) Survival analysis segregated by panel score, where a panel score > 6 indicates “nyesoPos=TRUE” and <6 indicates “nyesoPos=FALSE”.

**Figure S4.**
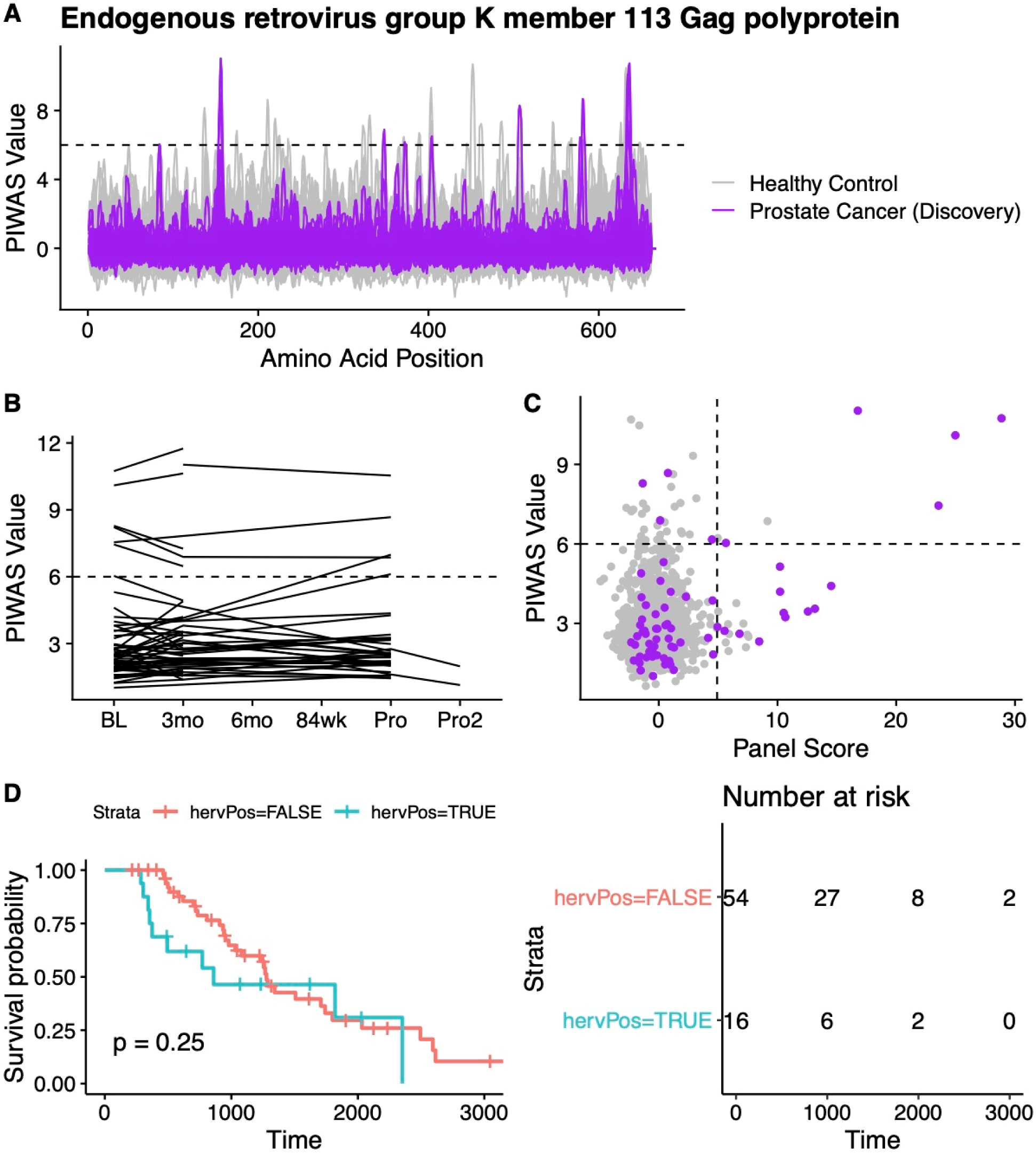
Additional findings for *HERVK-113*. (A) Complete tiling data for *HERVK-113* for every protein included in the PIWAS analysis. (B) PIWAS values across time for *HERVK-113*. One line is shown per patient. (C) Correlation between PIWAS value and panel score. (D) Survival analysis segregated by panel score, where a panel score > 6 indicates “nyesoPos=TRUE” and <6 indicates “nyesoPos=FALSE”.

